# Increased O-GlcNAcylation rapidly decreases GABA_A_R currents in hippocampus yet depresses neuronal output

**DOI:** 10.1101/672055

**Authors:** Luke T. Stewart, Kavitha Abiraman, John C. Chatham, Lori L. McMahon

**Author notes:** authors contributed equally (Luke T. Stewart, Kavitha Abiraman).

## Abstract

O-GlcNAcylation, a post-translational modification involving O-linkage of β-N-acetylglucosamine to Ser/Thr residues on target proteins, is increasingly recognized as a critical regulator of brain function in health and disease. Enzymes that catalyze O-GlcNAcylation are found at both presynaptic and postsynaptic sites, and O-GlcNAcylated proteins localize to synaptosomes. An acute increase in O-GlcNAcylation induces long-term depression (LTD) of excitatory transmission at hippocampal CA3-CA1 synapses, and depresses hyperexcitable circuits in vitro and in vivo. Yet, no study has investigated how O-GlcNAcylation modulates the efficacy of inhibitory neurotransmission. Here we show an acute increase in O-GlcNAc dampens GABAergic currents onto principal cells in rodent hippocampus likely through a postsynaptic mechanism, and has a variable effect on the excitation/inhibition balance. The overall effect of increased O-GlcNAc is reduced synaptically-driven spike probability via synaptic depression and decreased intrinsic excitability. Our results position O-GlcNAcylation as a novel regulator of the overall excitation/inhibition balance and neuronal output.

## Introduction

Synaptic integration and spike initiation in neurons is controlled by synaptic inhibition, which strongly influences neuronal output and information processing (Farrant and Nusser, 2005). Importantly, the balance of excitation to inhibition (E/I) is crucial to the proper functioning of circuits, and E/I imbalances have been implicated in a number of neuro-developmental disorders and neurodegenerative diseases including schizophrenia, autism spectrum disorders, and Alzheimer’s disease (Fernandez et al., 2007; Gogolla et al., 2009; Kehrer et al., 2008; Roberson et al., 2011). Thus, understanding the mechanisms that modulate the strength of inhibitory transmission is fundamental to unraveling how neuronal circuits function in normal and disease states.

Fast inhibitory transmission in the central nervous system is mediated by GABAA receptors (GABA_A_Rs), which are pentameric ligand-gated ion channels. The strength of this inhibition can be rapidly up- or down-regulated by post-translational modifications including ubiquitination (Saliba et al., 2007), palmitoylation (Keller et al., 2004; Rathenberg et al., 2004), and phosphorylation (Nakamura et al., 2015) of GABA_A_R subunits and/or associated proteins, which can alter channel function, trafficking, or stability at the membrane. While there is a vast body of literature on these post-translational modifications, O-GlcNAcylation, involving the O-linkage of β-N-acetylglucosamine (O-GlcNAc) to Ser/ Thr residues on target proteins, remains severely under-studied, with no examination to date of the effect of protein O-GlcNAcylation on inhibitory synapse physiology.

Addition and removal of O-GlcNAc are catalyzed by the single pair of enzymes O-GlcNAc transferase (OGT) and O-GlcNAcase (OGA), and is essential as genetic deletion of OGT and OGA are lethal (Keembiyehetty et al., 2015; Shafi et al., 2000). O-GlcNAcylation is metabolically-regulated and highly dynamic; global changes reversibly occur within minutes, and are dictated by availability of UDP-GlcNAc, which is synthesized from glucose via the hexosamine biosynthetic pathway (HBP) (Yang and Qian, 2017). The brain contains the second highest level of O-GlcNAcylated proteins in the body (Kreppel et al., 1997), with the hippocampus expressing one of the highest levels of OGT/OGA (Liu et al., 2004b). Notably, the majority of O-GlcNAcylated proteins are found at the synapse (Trinidad et al., 2012). However, only a handful of studies have examined the functional impact of O-GlcNAcylation in the brain (Lagerlöf et al., 2017; Hwang and Rhim, 2018; Ruan et al., 2014; Stewart et al., 2017; Taylor et al., 2014; Yang et al., 2017).

Previous work from our lab has examined the role of protein O-GlcNAcylation at glutamatergic synapses in hippocampal area CA1, where O-GlcNAcylation of the AMPA receptor (AMPAR) GluA2 subunit initiates long-term synaptic depression, termed ‘O-GlcNAc LTD’ (Taylor et al., 2014). Recent work (Hwang and Rhim, 2019) showing that increased O-GlcNAc suppresses excitatory transmission at CA3-CA1 synapses through the removal of GluA2 containing AMPARs is consistent with our findings. Additionally, increased O-GlcNAcylation dampens picrotoxin-induced epileptiform activity in area CA1 and CA3, and reduces seizure activity in the pentylenetetrazole in vivo model of seizure activity in mice (Stewart et al., 2017). Deletion of OGT in αCaMKII expressing neurons in the adult rodent brain causes a reduction in excitatory synaptic input onto hypothalamic PVN neurons (Lagerlöf et al., 2016), while OGT knock-out (KO) specifically from hypothalamic AgRP neurons reduces excitability via its effect on voltage-gated potassium channels (Ruan et al., 2014). Conversely, no studies to date have examined the role of protein O-GlcNAcylation in inhibitory synaptic function. Because serine phosphorylation of GABA_A_R subunits regulates the efficacy of neuronal inhibition (McDonald and Moss, 1997; Vithlani et al., 2011), it is highly likely that serine O-GlcNAcylation will also control GABA_A_R function.

Here, we show that an acute increase in protein O-GlcNAcylation rapidly induces a long-lasting decrease in strength of GABAergic synaptic transmission in hippocampus that is likely through an effect on post-synaptic GABA_A_Rs. This depression of inhibition produces a variable effect on the excitation/inhibition ratio in individual pyramidal cells, likely due to a simultaneous depression at excitatory synapses. However, the net effect in the intact circuit is a depression of neuronal output due to a simultaneous decrease in intrinsic excitability together with reduced synaptic drive at both excitatory and inhibitory synapses. Thus, global changes in O-GlcNAcylation induce complex changes in network activity by targeting excitatory and inhibitory synapses together with direct effects on intrinsic excitability.

## Results

### Acute increase in protein O-GlcNAcylation depresses GABAergic transmission onto CA1 pyramidal cells and dentate granule cells

To determine if protein O-GlcNAcylation modulates GAB-A^A^R-mediated inhibitory neurotransmission, we used whole-cell voltage clamp to record spontaneous inhibitory post-synaptic currents (sIPSCs) from CA1 pyramidal cells while blocking glutamatergic transmission using DNQX (10 μM) and DL-AP5 (50 μM). Following a 5 min baseline, we bath applied the HBP substrate glucosamine (GlcN, 5mM) and the OGA inhibitor thiamet-G (TMG, 1μM) to acutely increase protein O-GlcNAc levels, as done previously (Stewart et al., 2017; Taylor et al., 2014) (Fig 1Ai). We found a significant reduction in sIPSC amplitude (Fig 1Aii, cumulative probability distribution, p<0.0001, KS D value = 0.217, Kolmogorov-Smirnov test; inset: p<0.0001, Wilcoxon matched-pairs signed rank test) and inter-event interval (Fig 1Aiii, cumulative probability distribution, p<0.0001, KS D value = 0.084, Kolmogorov-Smirnov test; inset: p<0.0001, Wilcoxon matched-pairs signed rank test) in CA1 pyramidal cells. To ensure that the change in amplitude and inter-event interval of the sIPSCs was not a consequence of a technical artifact, sIPSCs during baseline and 5 min after GlcN+TMG application were averaged and scaled (Fig.1Aii and Bii, inset); the traces perfectly overlapped, indicating that the decrease in sIPSC amplitude and frequency observed are not due an increase in series resistance caused by prolonged recording or by washing on GlcN+TMG. It is also important to note that we observed a shift in holding current following application of GlcN+TMG (baseline: −138.2±13.6 pA vs. GlcN+TMG: −106.4±11.5 pA, n=9 cells, 5 rats, p=0.006, paired t-test), suggesting possible modulation of extrasynaptic GABA_A_Rs, which will be investigated in future experiments.

**Figure 1:**
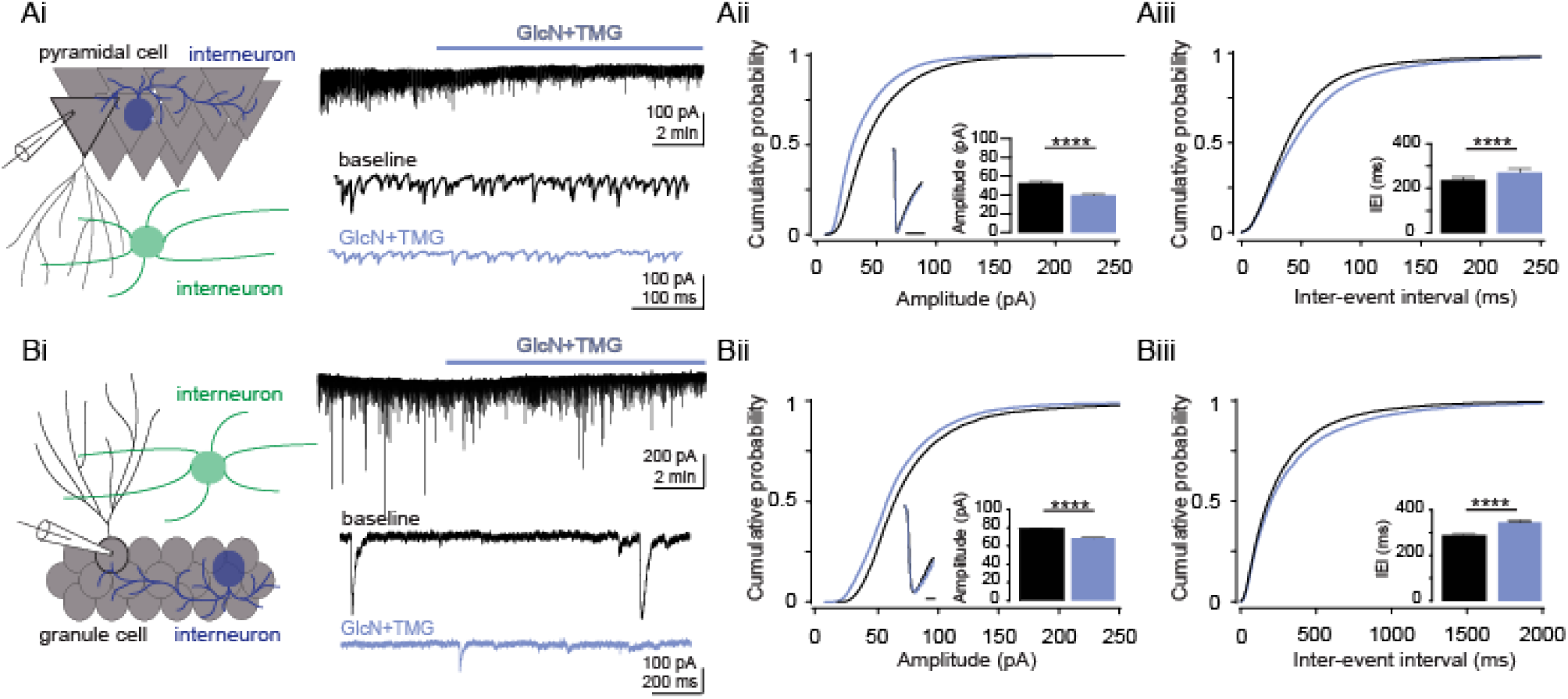
Acute increase in O-GlcNAcylation reduces spontaneous IPSCs in hippocampal principal cells. (Ai) (left) Schematic depicting recording set up in CA1. (right) representative sIPSC trace from CA1 pyramidal cell showing (top) GlcN+TMG wash on and (bottom) expanded time scale (control (black) and GlcN+TMG (blue)). (Aii) Cumulative probability distribution of sIPSC amplitude (p<0.0001, KS D value = 0.084, Kolmogorov-Smirnov test); inset: (left) scaled average sIPSC trace before (black) and after (blue) GlcN+TMG, scale bar: 5 ms. (right) average (±SEM) sIPSC amplitude. Baseline: 43.1±0.5pA, GlcN+TMG: 31.4±0.4 pA (p<0.0001, Wilcoxon matched-pairs signed rank test, n=9 cells, 5 rats).Inset shows no change in the rise-time or decay of averaged and scaled sIPSCs from before and after GlcN+TMG exposure. (Aiii) Cumulative probability distribution of sIPSC IEI p<0.0001, KS D value = 0.084, Kolmogorov-Smirnov test; inset: average (± SEM) sIPSC inter-event interval (IEI). Baseline: 53.1±0.9 ms, GlcN+TMG: 62.4±1.2 ms (p<0.0001, Wilcoxon matched-pairs signed rank test, n=9 cells, 5 rats). (Bi) (left) Schematic depicting recording set up in dentate gyrus and (right) representative sIPSC trace from a granule cell showing (top) GlcN+TMG wash on and (bottom) expanded time scale. (Bii) Cumulative probability distribution of sIPSC amplitude (p<0.0001, KS D value = 0.11, Kolmogorov-Smirnov test); inset: (left) scaled average sIPSC trace before (black) and after (blue) GlcN+TMG, scale bar: 5 ms. (right) average (± SEM) sIPSC amplitude. Baseline: 78.91±0.61pA, GlcN+TMG: 69.10±0.54 pA (p<0.0001, Wilcoxon matched-pairs signed rank test, n=11 cells, 7 rats).). Inset shows no change in the rise-time or decay of averaged and scaled sIPSCs from before and after GlcN+TMG exposure. (Biii) Cumulative probability distribution of sIPSC IEI; inset: average (± SEM) sIPSC IEI (p<0.0001, KS D value = 0.055, Kolmogorov-Smirnov test). Baseline: 301.4±3.6 ms, GlcN+TMG: 349.5±4.7 ms (p<0.0001, Wilcoxon matched-pairs signed rank test, n=11 cells, 7 rats). ****p<0.0001.

In order to determine if this decrease in sIPSCs induced by GlcN+TMG is specific to CA1 pyramidal cells or if it occurs at inhibitory synapses on other cells, we recorded sIPSCs from dentate gyrus granule cells (GCs) before and after bath application of GlcN+TMG (Fig 1Bi). We observed the same effect, namely a decrease in sIPSC amplitude (Fig 1 Bii, cumulative probability distribution, p<0.0001, KS D value = 0.109, Kolmogorov-Smirnov test; inset: p<0.0001, Wilcoxon matched-pairs signed rank test) and increase in inter-event interval (Fig 1Biii, cumulative probability distribution, p<0.0001, KS D value = 0.055, Kolmogorov-Smirnov test; inset: p<0.0001, Wilcoxon matched-pairs signed rank test), but no shift in holding current (p=0.768, paired t-test). These findings suggest that O-GlcNAc-mediated depression of GABAergic transmission is likely to be a general mechanism occurring at many inhibitory synapses in the brain.

The decrease in sIPSCs following increased O-GlcNAcylation could arise from a decrease in presynaptic GABA release, or from decreased postsynaptic GABA_A_R function. To distinguish between these possibilities, we recorded mIPSCs from both CA1 pyramidal cells and GCs in the presence of the voltage-gated sodium channel blocker TTX (1 μM) (Fig 2Ai and 2Bi). A change in mIPSC frequency typically reflects a change in presynaptic neurotransmitter release probability, while a change in mIPSC amplitude indicates a change in postsynaptic receptor function. Bath application of GlcN+TMG resulted in a reduction in mIPSC amplitude (Fig 2Aii, cumulative probability distribution, p<0.0001, KS D value = 0.24, Kolmogorov-Smirnov test; inset: p<0.0001, Wilcoxon matched-pairs signed rank test), but not inter-event interval (Fig 2Aiii, cumulative probability distribution, p=0.96, KS D value = 0.032, Kolmogor-ov-Smirnov test; inset: p<0.0001, Wilcoxon matched-pairs signed rank test) in CA1 pyramidal cells. We observed a similar decrease in mIPSC amplitude in dentate GCs (Fig 2Bii, cumulative probability distribution, p<0.0001, KS D value = 0.12, Kolmogorov-Smirnov test; inset: p<0.0001, Wilcoxon matched-pairs signed rank test), but in contrast to CA1, we also observe a slight but significant increase in mISPC inter-event interval (Fig 2Biii, cumulative probability distribution, p<0.0001, KS D value = 0.099, Kolmogor-ov-Smirnov test; inset: p<0.0001, Wilcoxon matched-pairs signed rank test). However, the very small difference in mean values between control (373.5±7.3 ms) and GlcN+T-MG (374.1±7.4 ms) suggests this significant difference may not be biologically relevant. Collectively, the decrease in mIPSC amplitude in the absence of a decrease in inter-event interval, at least in CA1 pyramical cells, is consistent with the interpretation that the dampening of inhibitory transmission following increased O-GlcNAcylation is due to a reduction in postsynaptic GABA_A_R function.

**Figure 2:**
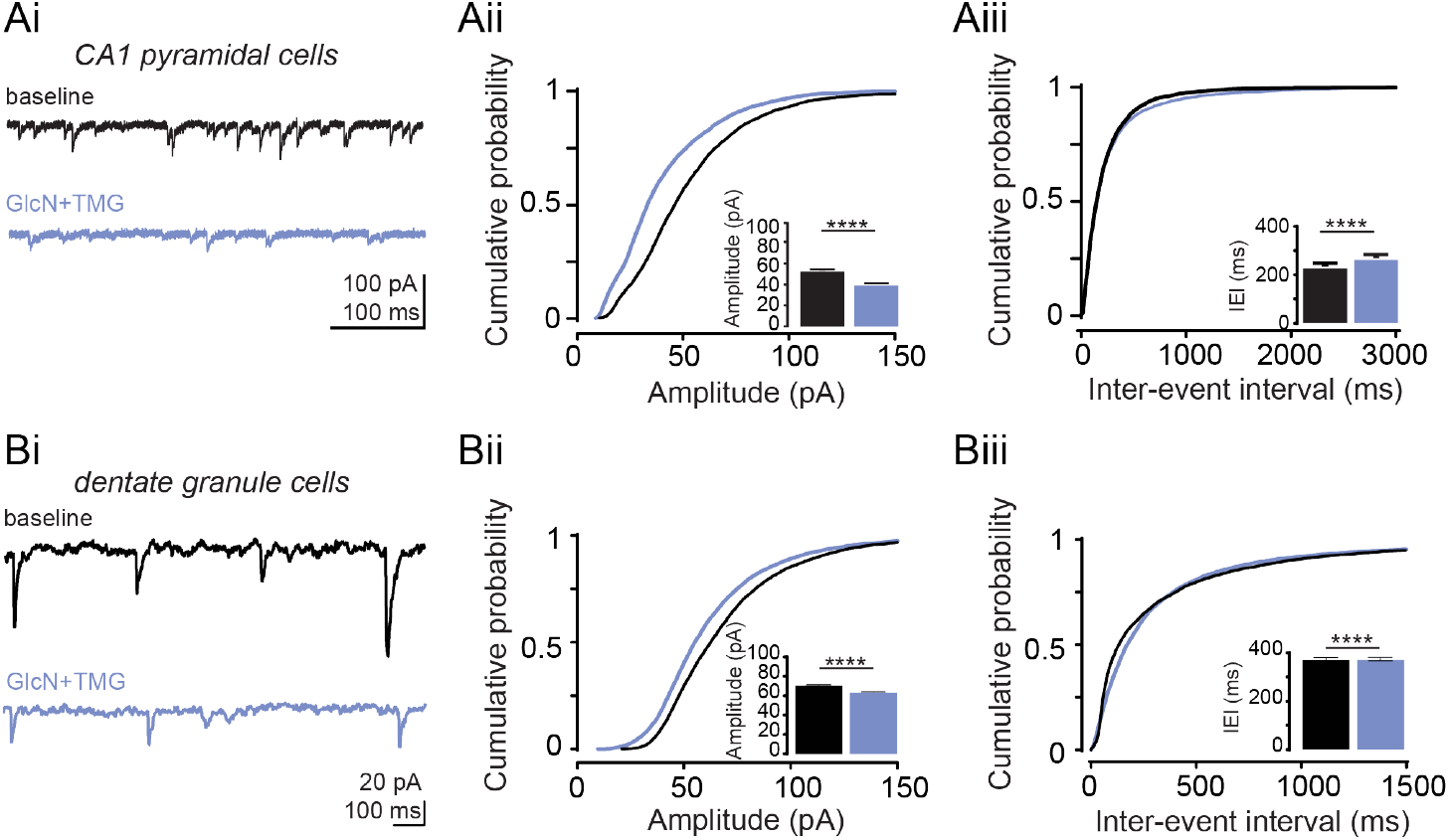
Acute increase in O-GlcNAcylation reduces miniature IPSC amplitude in hippocampal principal cells. (Ai) Representative mIPSC trace from CA1 pyramidal cell before (black) and after (blue) GlcN+TMG. (Aii) cumulative probability distribution of mIPSC amplitude (p<0.0001, KS D value = 0.24, Kolmogorov-Smirnov test); inset: average (± SEM) mIPSC amplitude. Baseline: 53.0±1.6 pA, GlcN+TMG: 40.0±1.0 pA (p<0.0001, Wilcoxon matched-pairs signed rank test, n=7 cells, 5 rats). (Aiii) Cumulative probability distribution of mIPSC IEI (p=0.96, KS D value = 0.032, Kolmogorov-Smirnov test); inset: average (± SEM) mIPSC IEI. Baseline: 236.7±14.3 ms vs GlcN+TMG: 271.5±16.2 ms (p=0.0005, Wilcoxon matched-pairs signed rank test, n=7 cells, 5 rats) (Bi) Representative mIPSC trace from dentate granule cell before (black) and after (blue) GlcN+TMG. (Bii) Cumulative probability distribution of mIPSC amplitude (p<0.0001, KS D value = 0.12, Kolmogorov-Smirnov test); inset: average (± SEM) mIPSC amplitude. Baseline: 71.1±0.4pA vs GlcN+TMG: 63.3±0.4 pA (p<0.0001, Wilcoxon matched-pairs signed rank test, n=12 cells, 6 rats). (Biii) Cumulative probability distribution of mIPSC IEI (p<0.0001, KS D value = 0.099, Kolmogorov-Smirnov test); inset: average (± SEM) mIPSC IEI. Baseline: 373.5±7.4 ms, GlcN+TMG: 368.8±7.5 ms (p=0.0005, Wilcoxon matched-pairs signed rank test, n=12 cells, 6 rats). ****p<0.0001.

### O-GlcNAcylation induces long-term depression of inhibitory transmission

We next tested if O-GlcNAcylation similarly affects the amplitude of electrically evoked IPSCs (eIPSCs), as evoked transmission may be dependent upon presynaptic vesicle pools (reviewed in Kavalali, 2015) distinct from those involved in spontaneous release. To address this and remaining questions, we focused our experiments on CA1 pyramidal cells. Following a 5 min stable baseline of eIPSCs (Cs Gluconate pipet solution; ECl-= −50 mV), bath applied GlcN+TMG elicited a significant reduction in eIPSC amplitude that lasted the duration of the recording (Fig 3 Ai and 3Aiii, p = 0.001, paired t-test), with no change in decay kinetics (Fig 3Aii, p=0,23, paired t-test).

**Figure 3:**
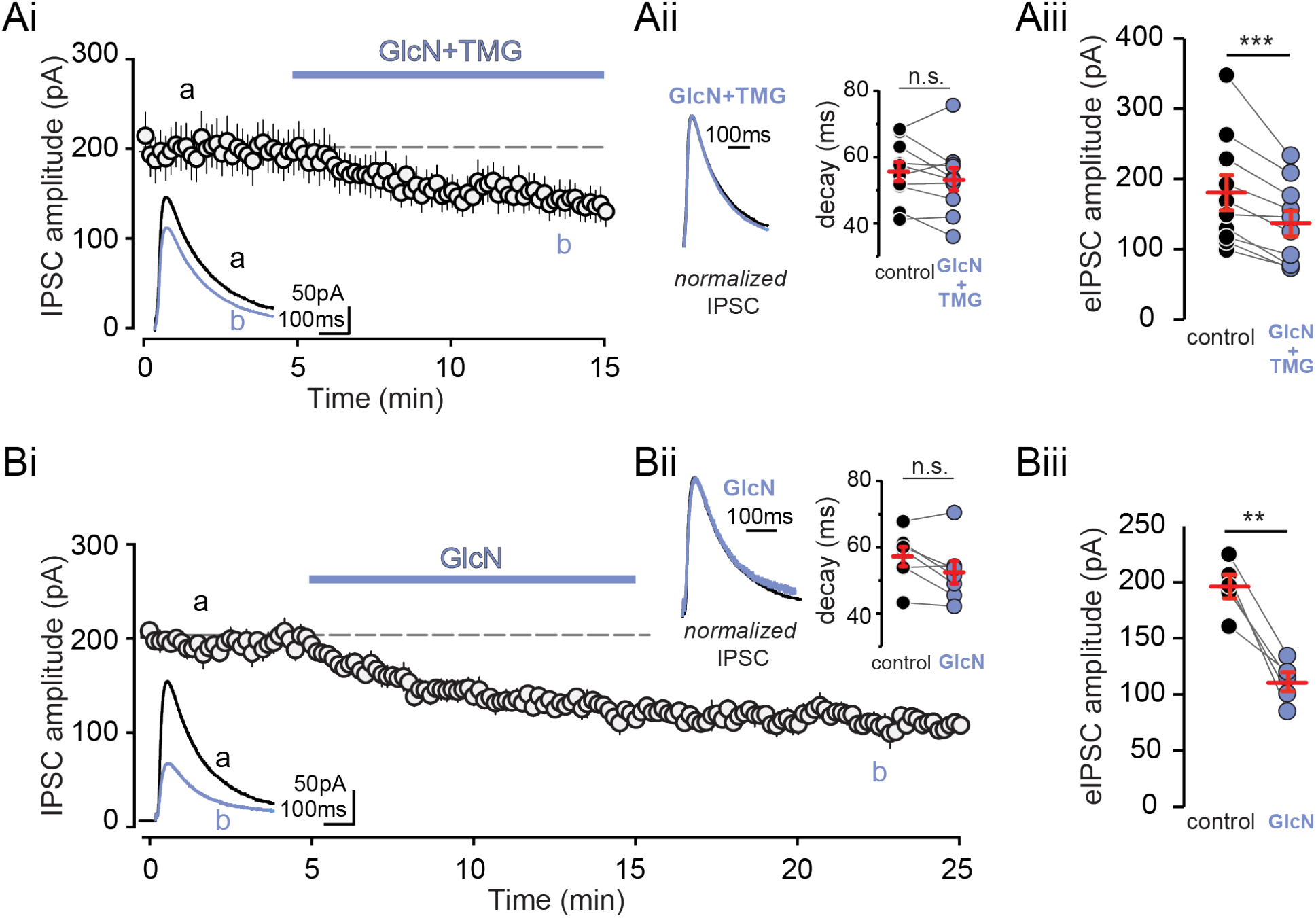
Increasing O-GlcNAcylation reduces evoked IPSC amplitude in CA1 pyramidal cells. (Ai) Group data showing average (±-SEM) evoked IPSC (elPSC) amplitude in control conditions and following GlcN+TMG wash on. Inset: representative elPSC traces before (black) and after GlcN+TMG (blue). (Aii) (left) Normalized representative eIPSC traces and (right) decay time before (black) and after GlcN+TMG (blue). Red horizontal bars represent the mean ± SEM. control: 55.7±2.9 ms vs GlcN+TMG: 53.4±3.4 ms (p=0.23, paired-test, n=10 cells, 5 rats). (Aiii) eIPSC amplitudes before (black) and after (blue) GlcN+TMG. Red horizontal bars represent the mean ± SEM. control: 180.9±25.1 pA vs GlcN+TMG 136.7±18.3pA (p=0.001, paired-test, n=10 cells, 5 rats). (Bi) Group data showing average (±SEM) evoked IPSC amplitude in control conditions and following GlcN wash on and wash out. Inset: representative eIPSC traces before (black) and after GlcN (blue) wash out. (Bii) (left) Normalized representative eIPSC traces and (right) decay time before (black) and after (blue) GlcN wash out. Red horizontal bars represent the mean ± SEM. control: 57.1±2.9 ms vs GlcN: 52.4±3.5 ms. (p=0.06, paired-test, n=5 cells, 2 rats). (Biii) eIPSC amplitudes before (black) and after (blue) GlcN wash out. Red horizontal bars represent the mean ± SEM. control: 196.3±10.5 pA vs GlcN+TMG 111.4±8.4 pA (p=0.003, paired-test, n=5 cells, 2 rats). **p<0.01,****p<0.001.

We previously reported that a 10 min exposure to GlcN (5mM) alone induces a significant increase in O-GlcNAcylation in hippocampus that is readily reversed within a 10 min washout, while O-GlcNAcylation remains elevated for up to 2 hrs following a 10 min exposure to TMG (Taylor et al., 2014). Importantly, the depression of transmission at CA3-CA1 synapses induced by GlcN outlasted the transient increase in protein O-GlcNAcylation, demonstrating expression of a long-term depression (LTD) at these synapses (Taylor et al., 2014). To test if O-GlcNAcylation similarly causes long-term depression at GABAergic synapses onto CA1 pyramidal cells, we applied GlcN alone for 10 min and recorded eIPSCs for 10 min during washout and observed a persistent reduction in amplitude (Fig 3Bi and 3Biii, p = 0.003, paired t-test), consistent with expression of a form of LTD at GABAergic synapses.

### O-GlcNAcylation has a variable effect on the E/I ratio

Since O-GlcNAcylation induces LTD of both excitatory (Taylor et al., 2014) and inhibitory transmission onto CA1 pyramidal cells (Figs.3), we wondered what the overall effect of increased O-GlcNAc is on the balance of excitation to inhibition (E/I). Thus, we measured the E/I ratio at CA3-CA1 synapses by recording compound synaptic currents consisting of mono-synaptic EPSCs followed by di-synaptic IPSCs in whole-cell recordings of CA1 pyramidal cells voltage-clamped at −30 mV. We observed a linear trend in the balance of E to I, with the E/I ratio increasing following GlcN+TMG application (Fig 4Ai, p<0.0001, repeated measures one-way ANOVA with post hoc test for linear trend). However, at the single cell level, the change in the E/I ratio was variable with just over half of cells (n=6/10) displaying a statistically increased ratio following GlcN+TMG exposure (p≤0.009, paired t-test on each individual cell comparing 5 mins before and 5 min following 5 min exposure to GlcN+TMG), and a statistically decreased E/I ratio in the rest (n=4/10; p≤0.01, paired t-test on each individual cell comparing 5 mins before and 5 min following 5 min exposure to GlcN+TMG). Thus, comparing the mean E/I ratios before and after GlcN+TMG exposure when all cells were averaged yields no significant difference (Fig 4Aii, p=0.08, paired t-test; red circles and blue circles indicate increased and decreased E/I respectively).

**Figure 4:**
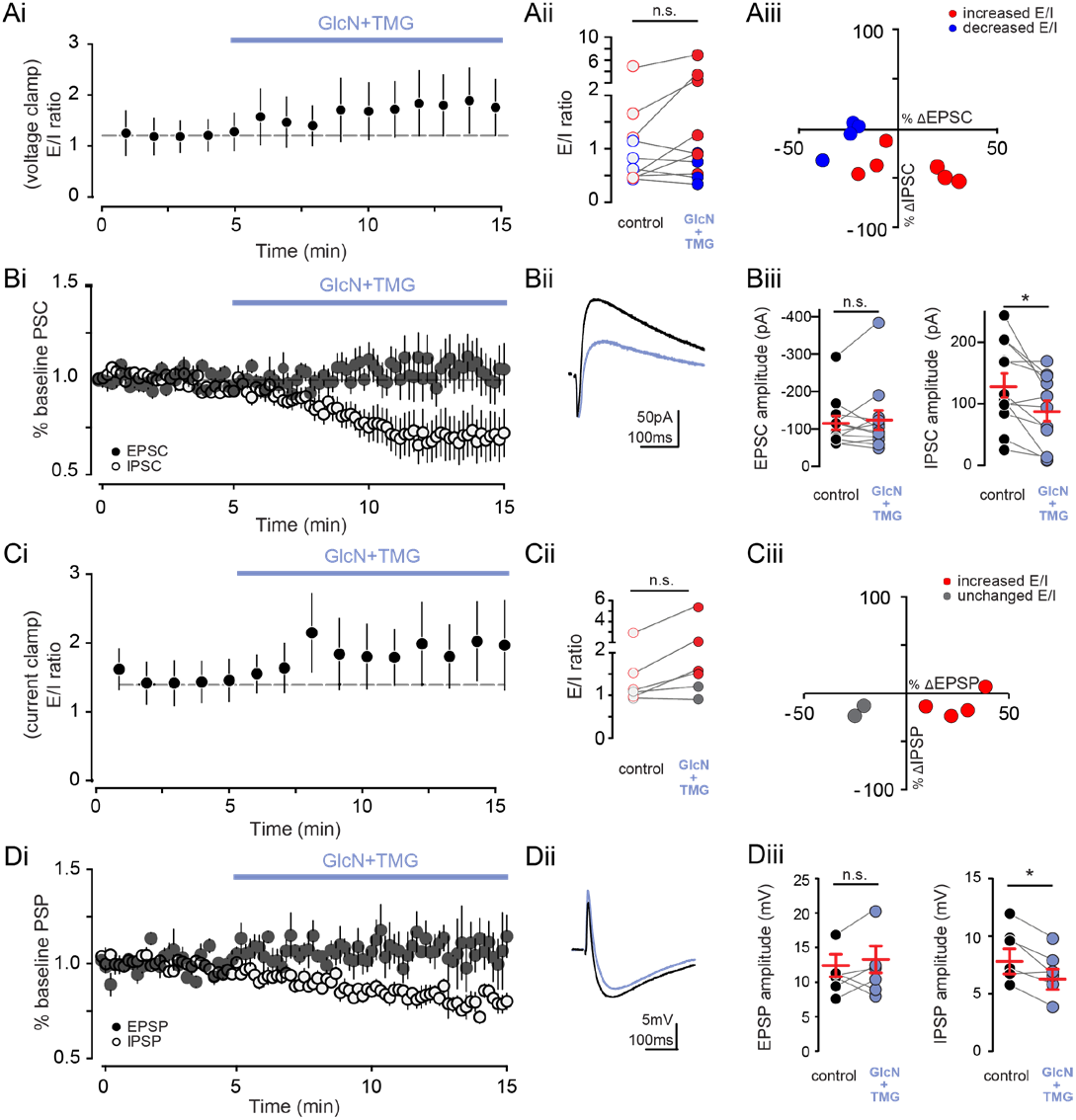
Increasing O-GlcNAcylation increases E-I ratio in pyramidal cells. (Ai) Group data showing average (±SEM) E/I ratio using voltage clamp in control conditions and following GlcN+TMG wash on (p<0.0001, repeated measures one-way ANOVA and post hoc test for linear trend, n=10 cells, 4 rats). (Aii) E/I ratio before (open circles) and after (filled circles) GlcN+TMG. control: 1.2±0.4 vs GlcN+TMG: 1.8±0.6 (p=0.08, paired t-test, n=10 cells, 4 rats). Red circles indicate increased E/I and blue circles indicate decreased E/I ratio. (Aiii) Relationship between percent change in the EPSC and IPSC following GlcN+TMG from the recordings in Ai. Red circles indicate increased E/I and blue circles indicate decreased E/I ratio. (Bi) Normalized group data showing average (±-SEM) evoked mono-synaptic EPSC (filled circles) and di-synaptic IPSC (open circles) amplitude in control conditions and following GlcN+TMG wash on.(Bii) Representative compound EPSC and IPSC before (black) and after (blue) GlcN+TMG. (Biii) EPSC amplitude before (black) and after (blue) GlcN+TMG. Red horizontal bars represent the mean ± SEM. control: −120.2±22.1 pA vs GlcN+TMG: −116.1±30.7 pA (p=0.43, Wilcoxon matched-pairs signed rank test, n=10 cells, 4 rats). (right) IPSC amplitude before (black) and after

We next examined the compound EPSCs and IPSCs individually as both can be depressed by inceased O-GlcNAc. GlcN+TMG application caused a significant reduction in the IPSC component of the compound current (Fig 4Bi-Biii, open circles shown as %baseline current, paired t-test, p = 0.03), with no significant change in the EPSC component (Fig 4Bi-Biii, closed circles shown as % baseline current, p=0.43, Wilcoxon matched-pairs signed rank test). These data indicate that on average, increasing O-GlcNAc elicits a greater depression of the disynaptic GABA_A_R-mediated IPSC than the mono-synaptic glutamatergic EPSC. Normally, GABA_A_R-mediated transmission limits the amplitude of EPCSs through shunting the excitatory potential, and when the inhibitory shunt is decreased or blocked, the EPSC amplitude increases. Thus, decreasing the inhibitory shunt via the O-GlcNAc-mediated decrease in IPSC amplitude predicts that the EPSC amplitude should increase (Mitchell and Silver, 2003), which we observed in a fraction of the cells (n=3/10, Fig 4Aiii). On the other hand, the GluA2-AMPAR mediated O-GlcNAc LTD at CA3-CA1 synapses (Taylor et al., 2014), occurring in parallel with the decrease in IPSC amplitude could occlude the expected increase in EPSC amplitude cause by the decreased inhibitory shunt, which we observed in the majority of our cells (n=7/10, Fig 4Aiii). Thus, the overall effect of increasing O-GlcNAc is extremely complicated given that exposure to GlcN+TMG causes synaptic depression at both glutamatergic and GABAergic synapses within minutes, and the impact on the E/I ratio will be driven by the magnitude of the change in the EPSC vs IPSC, with increases in the E/I ratio being driven by a larger increase in the EPSC or smaller decrease in the EPSC amplitude compared to the IPSC (compare blue and red circles in Fig 4Aiii). We also confirmed this variable effect on the E/I ratio in current clamp recordings from CA1 pyramidal cells, and found, once again, a linear trend for an increase in the E/I ratio (Fig 4Ci;, p<0.01, repeated measures one-way ANOVA with post hoc test for linear trend) following GlcN+TMG exposure, and a greater depression of the IPSP vs EPSP (Fig 4Di-Diii, IPSP vs EPSP: p=0.01 and p=0.4, paired t-test) translating into an increase in the E/I ratio in the majority of cells (Fig 4Cii, n=4/6 cells, p≤0.01, paired t-test on each individual cell comparing 5 mins before and 5 min following 5 min exposure to GlcN+TMG).

### O-GlcNAcylation reduces action potential probability in CA1 pyramidal cells

How does this variable effect of increased O-GlcNAc on the E/I ratio affect neuronal output? In a previous study, we reported that increasing O-GlcNAc decreased basal spontaneous activity of CA3 pyramidal cells in an intact circuit, and depressed epileptic activity in area CA1 in hyperexcitable conditions generated by blocking GABA_A_Rs (Stewart et al., 2017). Therefore, to determine the net effect of increasing protein O-GlcNAcylation on CA1 pyramidal cell output when the circuit is intact, we carried out current-clamp recordings from CA1 pyramidal cells and measured synaptically driven AP probability before and after application of GlcN+TMG. Stimulus intensity was set to achieve a ~25-50% success rate of generating a synaptically driven AP during the 5min baseline (Fig 5Ai, 0-5min) after which GlcN+TMG were bath applied to increase O-GlcNAc levels (Fig 5Ai, 5-20min). Increasing O-GlcNAcylation caused a significant reduction in AP probability in all recorded cells, despite the increase in E/I ratio (Fig 5Aii and Aiii, p < 0.0001, paired t-test). Importantly, control experiments were interleaved to rule out technical artifacts since AP probability is particularly sensitive to any change in the quality of the recording (Fig 5Bi-Biii, p = 0.56, paired t-test).

**Figure 5:**
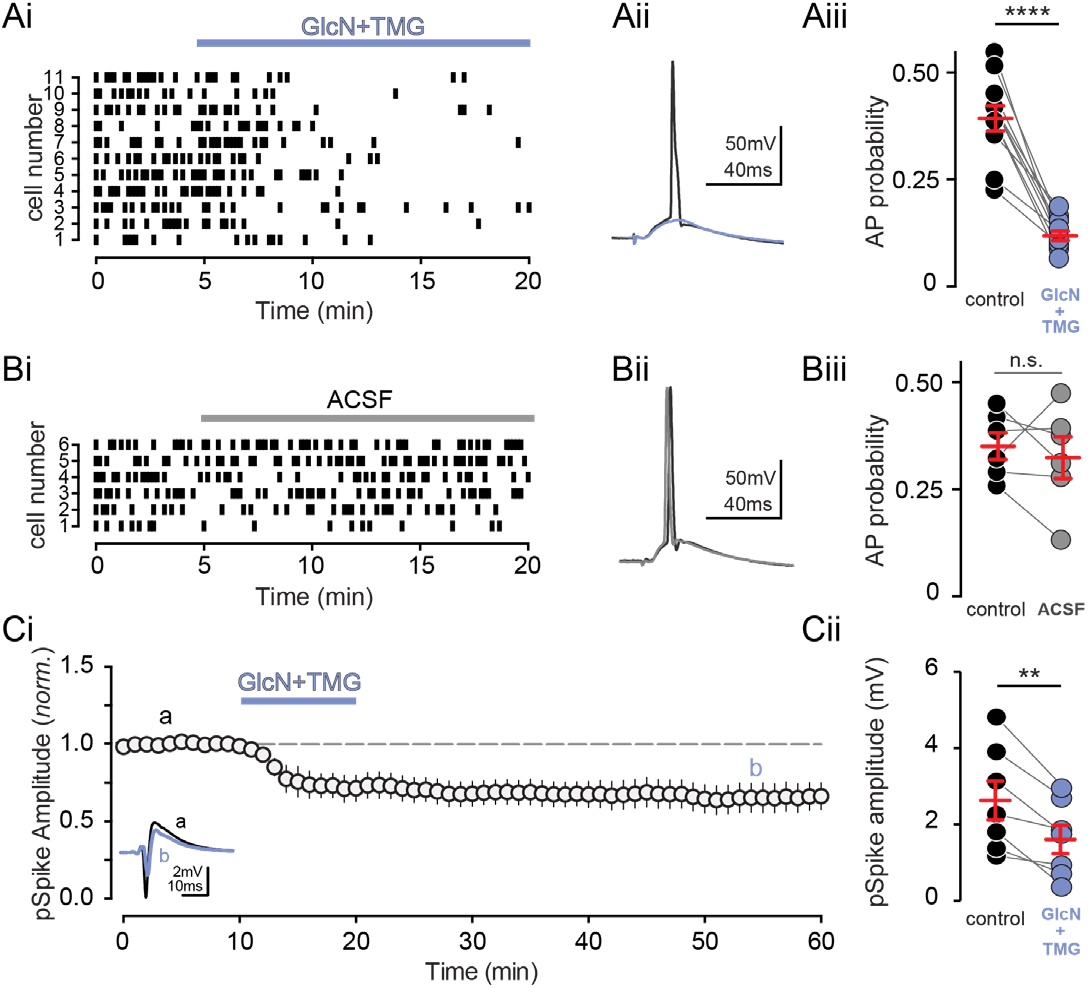
O-GlcNAcylation reduces CA1 action potential output. (Ai) Raster plot and (Aii) representative trace of synaptically-evoked APs before (black) and after GlcN+TMG (blue). (Aiii) AP probability before (black) and after (blue) GlcN+TMG. Red bars represent mean± SE. control: 0.39±0.03 vs. GlcN+TMG: 0.12±0.01 (p<0.0001, paired t-test, n=11 cells, 6 rats). (Bi) Raster plot and (Bii) representative average of synaptically-evoked APs before (black) and after (grey) ACSF application as a negative control. (Biii) Quantification of AP probability before (black) and after (grey) ACSF. Red bars represent mean ± SE. control: 0.35±0.03 vs. GlcN+TMG: 0.33±0.05 (p=0.5622, paired t-test, n=6 cells, 4 rats). (Ci) Normalized Group data showing pop-spike amplitude in baseline conditions and after GlcN+TMG (left). inset: representative trace showing pop spike before (black) and after (blue) GlcN+TMG. (Cii) pSpike amplitude before (black) and after (blue) GlcN+TMG. Red bars represent mean ± SEM. control: 2.64±0.52 mV vs. GlcN+TMG: 1.61 ±0.38 mV (p=0.004, paired t-test, n=7 slices, 3 rats).**p<0.01, ****p<0.0001.

As a further test of the effects of increasing O-GlcNAcylation on the net activity in the intact circuit, we performed a complementary experiment measuring the output of the CA1 population by recording extracellular population spikes (pSpike) in the CA1 pyramidal cell layer during a 10 min application of GlcN+TMG (Fig 5Ci, 10-15min). Consistent with the single-cell recordings, we observed a significant and sustained reduction in pSpike amplitude following an increase in O-GlcNAcylation (Fig 5Cii, p = 0.004, paired t-test), which supports our previous report of decreased CA3 output in the context of increased protein O-GlcNAcylation (Stewart et al., 2017).

### Reduced intrinsic excitability of CA1 pyramidal cells contributes to the O-GlcNAc-mediated depression of excitability

The decrease in synaptically-evoked AP probability following increased O-GlcNAcylation may arise as a consequence of non-mutually exclusive synaptic and intrinsic mechanisms. Therefore, we next tested the possibility that O-GlcNAcylation decreases intrinsic excitability by modulating the active and passive properties of CA1 pyramidal cells. We used whole-cell current clamp recordings of CA1 pyramidal cells during hyperpolarizing and depolarizing current injections in the presence of pharmacological blockers of GABA_A_Rs, AMPARs, and NMDARs, to isolate the cells from synaptic inputs (Fig 6A). We found that O-GlcNAcylation caused a significant reduction in the input resistance (Fig 6C, p=0.031, paired t-test) when cells were isolated from the synaptic network, and a significant increase in rheobase, or the minimum current required to elicit a single action potential (Fig 6D, p=0.036, paired t-test). We also observed an increase in AP threshold (Fig 6E, p=0.030, paired t-test), but no difference in AP shape (Fig 6F, 6G, 6H, AP amplitude: p=0.441; AP rise: p=0.143; AP half width: p=0.653, paired t-test). Additionally, we found a small but significant decrease in the number of action potentials generated with higher current injection steps following GlcN+TMG application (FO-GlcNAc(1,8)= 5.81, p<0.05, two-way repeated measures ANOVA). These changes are consistent with a decrease in intrinsic excitability.

**Figure 6:**
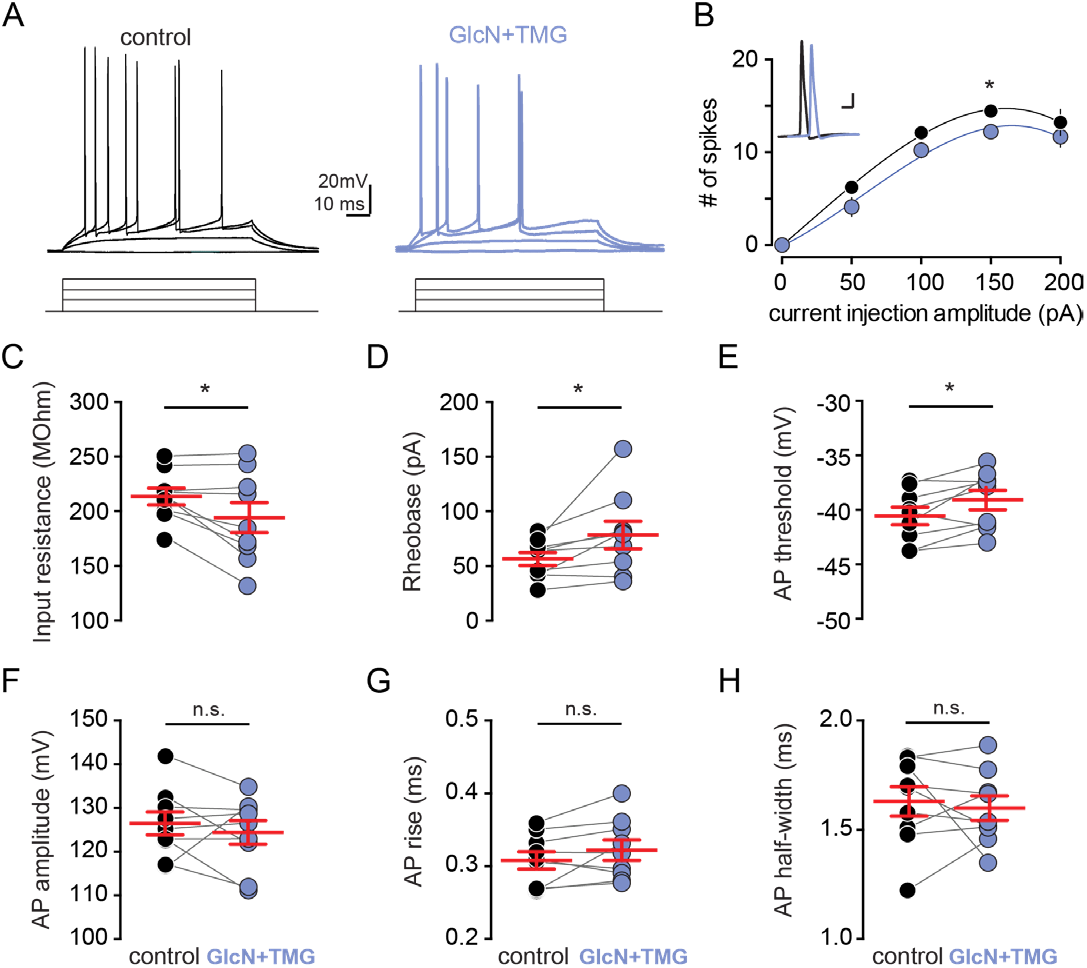
O-GlcNAcylation reduces intrinsic excitability of CA1 pyramidal cells. (A) Voltage responses to current injections (bottom, 800 ms, 20 pA steps) before (black) and after (blue) GlcN+TMG. (B) Number of spikes elicited by increasing current steps before (black) and after (blue) GlcN+TMG. Circles represent mean ± SEM. Inset: AP waveform before (black) and after (blue) GlcN+TMG. (scale bar: 20 mV, 10 ms). (C) Input resistance, (D) rheobase, (E) AP threshold, (F) AP amplitude, (G) AP rise, and (H) AP half-width before (black) and after (blue) GlcN+TMG. Red bars indicate mean ± SEM. Input resistance: 213.4 ± 7.7 MΩ (control) vs 194.1± 13.7 MΩ (GlcN+TMG) (p=0.031, paired t-test). Rheobase: 56.7±5.9 pA (control) vs 78.4±12.5 pA (GlcN+TMG) (p=0.036, paired t-test). AP threshold: −40.6±0.8 mV (control) vs −39.1±0.9 mV (GlcN+TMG) (p=0.030, paired t-test). AP amplitude: 126.3±2.6 mV(control) vs 124.3±2.7 mV (GlcN+TMG) (p=0.44, paired t-test). Rise: 0.31±0.01 ms (control) vs 0.32±0.01 ms (GlcN+TMG) (p=0.143, paired t-test). AP half-width: 1.63± 0.07 ms(control) vs 1.59± 0.06 (GlcN+TMG) (p=0.653, paired t-test, n= 9 cells, 2 rats). *p<0.05.

Collectively, our current and previously published data (Stewart et al., 2017) suggest that both synaptic and intrinsic mechanisms contribute to a net decrease in CA1 pyramidal cell excitability, despite a larger O-GlcNAc-mediated depression of inhibitory versus excitatory transmission. Therefore, to distinguish the decreased intrinsic excitability from the synaptic depression at excitatory synapses, we repeated the pSpike recordings in GluA2 KO mice in which O-GlcNAc LTD at CA3-CA1 synapses will be absent (Fig 7Ai and 7Aii). Interestingly, increasing O-GlcNAcylation depressed pSpike amplitude in both the wild-type and GluA2 KO mice (Fig 7Aiii, control: p<0.01 KO: p<0.01, sidak’s multiple comparisons test), while the persistent depression (O-GlcNAc LTD) following GlcN+TMG wash-out was only observed in wild-type mice, as expected (Fig 7Aiii, control: p<0.01 KO: p>0.05, sidak’s multiple comparison test). Thus, the early depression of the pSpike amplitude during GlcN+TMG application is likely a consequence of decreasing CA1 pyramidal cell intrinsic excitability, while the persistent depression of excitability is a consequence of synaptic depression mediated by O-GlcNAc LTD.

**Figure 7:**
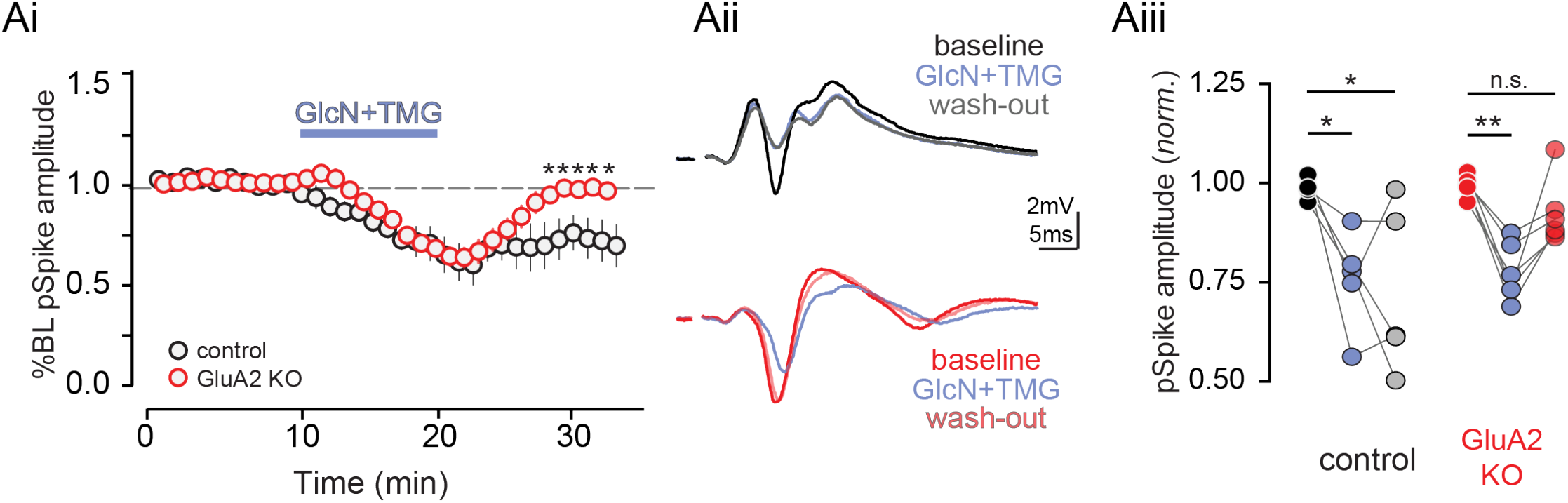
Increased O-GlcNAcylation reduces CA1 spike output. (Ai) Normalized group data showing pop-spike amplitude in baseline conditions and after GlcN+TMG in control (black) and GluA2 KO (red) mice (*p<0.05, two-way ANOVA with Sidak’s multiple comparison test, n= 5 slices, 2 rats). (Aii) Representative averaged traces showing (top) pop spike amplitude before (black), after (blue) GlcN+TMG, and following wash-out (grey) in control mice and (bottom) pop spike amplitude before (red), after (blue) GlcN+TMG and following wash-out (pink) in GluA2 KO mice. (Aiii) Pop spike amplitude before (black), after (blue) GlcN+TMG and following wash-out (grey) in control mice (baseline vs. GlcN+TMG: p<0.05; baseline vs. wash-out: p<0.05, repeated measures one-way ANOVA with Sidak’s multiple comparison’s test, n= 5 slices, 2 rats) and pop spike amplitude before (red), after (blue) GlcN+TMG, and following wash-out (pink) in GluA2 KO mice (baseline vs. GlcN+TMG: p<0.01; baseline vs. wash-out: p>0.05, repeated measures one-way ANOVA with Sidak’s multiple comparison’s test, n= 5 slices, 2 rats). *p<0.05, **p<0.01.

## Discussion

We provide, for the first time, empirical evidence of a role for protein O-GlcNAcylation in regulating synaptic inhibition in principal cells of the hippocampus. First, we show that acutely increasing O-GlcNAcylation reduces the frequency and magnitude of sIPSCs in CA1 pyramidal cells and dentate granule cells, and selectively decreases the amplitude of mIPSCs, suggesting a postsynaptic mechanism. The amplitude of evoked IPSCs onto CA1 pyramidal cells is similarly reduced, and the depression is long-lasting, suggesting a form of LTD induced by O-GlcNAc at GABAergic synapses. Second, in a hippocampal circuit with intact excitation and inhibition, an acute increase in O-GlcNAcylation has a variable effect on the E/I ratio, despite a greater effect on inhibition vs excitation. Finally, we show that O-GlcNAcylation modulates the final output of CA1 pyramidal cells by reducing the action potential probability through intrinsic and synaptic mechanisms. These results position protein O-GlcNAcylation as a potent regulator of both synaptic inhibition as well as excitation, as we reported previously (Stewart et al., 2017; Taylor et al., 2014).

While the effect of the analogous post translational modification, phosphorylation, on synaptic transmission has been extensively characterized, little is known about the role of O-GlcNAcylation on excitatory and inhibitory transmission. We have previously shown that acute increases in O-GlcNAcylation causes a GluA2 AMPAR subunit dependent long-term depression of excitatory transmission, O-GlcNAc LTD, in CA1 pyramidal cell dendritic field potentials and CA1 population spikes (Stewart et al., 2017; Taylor et al., 2014). Here, we report that an acute increase in protein O-GlcNAcylation similarly causes a depression of inhibition onto CA1 pyramidal cells and dentate granule cells. This includes a reduction in the amplitude but not frequency of mIPSCs onto hippocampal principal cells, suggesting a post-synaptic site of action. This reduction in inhibition could involve a number of mechanisms including internalization of GABA_A_Rs, a change in the conductance of individual GABAARs, or destabilization of scaffolding proteins that anchor GABA_A_Rs to the membrane. Indeed, recent work (Hwang and Rhim, 2019) suggests the reduction in mEPSC amplitude following increased O-GlcNAcylation involves the endocytosis of GluA2 subunit containing AMPARs, which likely explains expression of the GluA2-dependent LTD we previously reported (Taylor et al., 2014). Additionally, phosphorylation of specific serine/threonine residues on GABA_A_Rs modulates inhibitory transmission, an effect that includes a reduction in GABA_A_R currents (Nakamura et al., 2015), which might also occur following increased O-GlcNAc.

Notably, phosphorylation alters inhibition in the brain in a kinase- and region-specific manner (for review, see Nakamura et al., 2015)). Activation of PKA depresses synaptic GABA_A_R function in CA1 pyramidal cells but not dentate gyrus granule cells (GCs), whereas PKC activation alters mini-IPSC amplitude in GCs alone (Poisbeau et al., 1999). O-GlcNAcylation, however, is uniquely positioned to turn down global inhibition in a non region-specific manner as only one enzyme, OGT, is known to catalyze its addition to proteins. While phosphorylation and O-GlcNAcylation on some proteins can occlude one another (Cheng et al., 2000; Chou et al., 1995; Dias et al., 2009; Griffith and Schmitz, 1999; Liu et al., 2004a), other times they can act independently (Taylor et al., 2014) or synergistically (for review, see Zeidan and Hart, 2010). Whether there is a negative or positive interplay between the effects of phosphorylation and O-GlcNAcylation on inhibition remains to be determined.

We show that acute increases in O-GlcNAcylation rapidly depress the di-synaptic IPSC while having a variable effect on the amplitude of the EPSC. This variability is likely explained by competing mechanisms including an O-GlcNAc mediated decrease in the inhibitory shunt and O-GlcNAc LTD at excitatory synapses. These changes are complex as indicated by the variable effect on the E/I ratio with some cells experiencing a significant increase and others a decrease such that a significant change in the averaged dataset is not observed. Despite this, there is a consistent reduction in the synaptically-evoked action potential probability in pyramidal cells. While subthreshold EPSPs can spatially and temporally summate to generate APs, the efficacy of EPSP to spike coupling is thought to depend on several factors including IPSP magnitude and timing (Pouille and Scanziani, 2001), as well as passive and active properties of the post-synaptic neuron (Magee, 2000). Interestingly, (i) individual pyramidal cells vary widely in their threshold afferent stimulation required for spiking, but this threshold is controlled by the magnitude of the EPSC and not the amplitude or timing of inhibition (Pouille et al., 2009); and (ii) increasing O-GlcNAcylation leads to a depression of field EPSPs in area CA1, likely owing to an internalization of GluA2 containing AMPARs (Hwang and Rhim, 2019; Stewart et al., 2017). Thus, it stands to reason that the decrease in AP probability could stem from a concomitant depression of the CA3-CA1 EPSC. We found that O-GlcNAcylation not only reduces the AMPAR mediated dendritic depolarization (Taylor et al., 2014), but also changes the intrinsic properties of CA1 neurons, making it harder for the cell to spike. This change in intrinsic properties following an increase in O-GlcNAc is consistent with recently published work by Hwang and Rhim, 2019 showing a reduction in depolarizing voltage-gated sodium channel currents and an increase in hyperpolarizing voltage-gated potassium currents. We have also previously reported a reduction in basal spontaneous spiking activity in area CA3 of hippocampus following increased O-GlcNAc (Stewart et al., 2017), further supporting a suppressive effect of O-GlcNAcylation on neuronal output. Notably, this finding is contrary to studies using genetic strategies to increase O-GlcNAcylation (Yang et al., 2017), which reported no changes in intrinsic excitability, or mini-EPSCs and -IPSCs following heterozygous loss of function mutation of OGA, whose protein product catalyzes the removal of O-GlcNAc from proteins. The conflicting results are mostly likely due to differential effects of acute versus chronic upregulation in protein O-GlcNAcylation, as chronic changes can produce compensatory mechanisms to maintain synaptic function.

Studying the effect of acute, moment-to-moment changes in global O-GlcNAc levels on synaptic transmission likely reflects the impact of metabolically driven changes in O-GlcNAcylation occurring in vivo under both physiological and pathophysiological conditions. The OGT substrate UDP-GlcNAc is produced by the flux of glucose through the hexosamine biosynthetic pathway, which incorporates nucleotides, fatty acids, and amino acids. As such, O-GlcNAcylation is positioned to operate as a nutrient sensor (for review, see Lagerlöf, 2018). For instance, fasting promotes energy conservation via the upregulation of OGT and O-GlcNAc and its effect on the activity of hypothalamic neurons (Ruan et al., 2014). Conversely, chronic elevation of glucose can also increase O-GlcNAcylation of certain targets, contributing to cardiac and neuronal pathophysiology (Erickson et al., 2013). Additionally, O-GlcNAcylation plays a crucial role in pathological states such as diabetes (Erickson et al., 2013; Vaidyanathan and Wells, 2014) where its elevation can contribute to disease pathogenesis, in epilepsy where increasing O-GlcNAc is protective (Sánchez et al., 2019; Stewart et al., 2017), and in neurodegeneration (Levine et al., 2019; Wang et al., 2016; Yuzwa and Vocadlo, 2014; Yuzwa et al., 2008, 2012, 2014a, 2014b; Zhu et al., 2014), where increasing O-GlcNAc can decrease pathological accumulation of phosphorylated tau or α-synuclein. Collectively, studies published by us and others suggest that maintaining proper balance in O-GlcNAcylation is critical to maintaining normal hippocampal function, as too much or too little O-GlcNAc has been linked to deficits in learning and memory (Taylor et al 2014; Yang et al 2017; Wang et al 2016). Revealing the complex changes in neuronal and synaptic function induced by alterations in O-GlcNAcylation in non-pathological and pathological states will require further investigation.

## Materials and methods

All experimental procedures were approved by the University of Alabama at Birmingham Institutional Animal Care and Use Committee and follow the National Institutes of Health experimental guidelines.

Hippocampal slice preparation. Male and female Sprague Dawley rats (age 4-8 weeks; Charles River Laboratories) or male and female mice (age 4 – 12 weeks, GluA2 KO, Jax Labs #002913) were anesthetized with isoflurane, rapidly decapitated, and brains removed; 400μm (rats) or 350 m (mice) coronal slices from dorsal hippocampus were prepared on a VT1000P vibratome (Leica Biosystems) in oxygenated (95%O2/5%CO2) ice-cold, high sucrose cutting solution (in mM as follows: 85.0 NaCl, 2.5 KCl, 4.0 MgSO4, 0.5 CaCl2, 1.25 NaPO4, 25.0 glucose, 75.0 sucrose). After cutting, slices were held at room temperature for 1 to 5 hr in a submersion chamber with continuously oxygenated standard ACSF (in mM as follows: 119.0 NaCl, 2.5 KCl, 1.3 MgSO4, 2.5 CaCl2, 1.0 NaH2PO4, 26.0 NaHCO3, 11.0 glucose) with 2mM kynurenic acid to better preserve slice health.

Electrophysiology. All recordings were performed in a submersion chamber with continuous perfusion of oxygenated standard ACSF. The blind patch technique was used to acquire whole-cell recordings from CA1 pyramidal neurons and dentate granule cells (GCs). Neuronal activity was recorded using an Axopatch 200B amplifier and pClamp10 acquisition software (Molecular Devices, Sunnyvale, CA). Signals were filtered at 5 kHz and digitized at 10 kHz (Digidata 1440). Patch pipettes (BF150-086 or BF150-110; Sutter Instruments, Novato, CA) were pulled on a Sutter P-97 (Sutter Instruments, Novato, CA) horizontal puller to a resistance between 2-6 MΩ. Spontaneous inhibitory post-synaptic currents (sIPSCs) were pharmacologically isolated with bath perfusion of DNQX (10μM; Sigma) and DL-AP5 (50μM; Tocris) or R-CPP (5 μM; Abcam). Purity of GABA_A_R currents was verified with perfusion of picrotoxin (50μM; Sigma) following experimental recording. Spontaneous and miniature IPSCs were recorded using CsCl internal solution (in mM: 140.0 CsCl, 10.0 EGTA, 5.0 MgCl2, 2.0 Na-ATP, 0.3 Na-GTP, 10.0 HEPES; ECl = 0mV). Evoked GABAAR currents were recorded using Cs-gluconate internal solution (in mM: 100.0 Cs-gluconate, 0.6 EGTA, 5.0 MgCl2, 2.0 Na-ATP, 0.3 Na-GTP, 40.0 HEPES; ECl = −61.5 mV) with a twisted nichrome wire bipolar electrode positioned in stratum radiatum (100μs, 0.1Hz). Whole-cell current clamp recordings from CA1 pyramidal neurons were carried out using K-gluconate internal solution (in mM: 120.0 K-gluconate, 0.6 EGTA, 5.0 MgCl2, 2.0 Na-ATP, 0.3 Na-GTP, 20.0 HEPES; ECl = −61.5 mV). All cells were dialized for 3-7min prior to the beginning of experimental recordings. Stability of series resistance was verified using post-hoc scaling of averaged waveforms before and after pharmacologicaly increasing O-GlcNAcylation. pSpikes in the CA1 cell body layer were evoked by stimulating Schaffer collaterals with pairs of pulses (0.1 Hz, 100 μs duration at 50 ms interval) in stratum radiatum with a twisted nichrome wire bipolar electrode and recorded with a glass pipet filled with aCSF placed nearby in stratum pyramidale.

Modulation of O-GlcNAc levels. As previously described (Taylor et al., 2014; Stewart et al., 2017), O-GlcNAcylation was acutely increased via bath application of glucosamine (GlcN, 5 mM; Sigma) alone or in combination with the selective OGA inhibitor TMG (1 μM; Chem Molecules), the most potent OGA inhibitor (Ki = 21 nM; human OGA) currently available (Macauley et al., 2005; Yuzwa et al., 2008). GlcN and TMG were used in combination to ensure robust and lasting increases in protein O-GlcNAc levels. In some experiments, we tested whether applying the HBP substrate GlcN alone was sufficient to modulate inhibitory transmission, as this approach more closely recapitulates the endogenous regulation of cellular O-GlcNAcylation.

Experimental design and statistical analysis. Recordings were analyzed using Clampfit 10.6. Outliers were determined as per outlier test in GraphPad Prism 8.1.0, La Jolla, CA, and excluded. For two-group comparisons, statistical significance was determined by two-tailed paired or unpaired Student’s t-tests (parametric), or Wilcoxon matched-pairs signed rank test (non-parametric, paired). Multi-groups were analyzed using repeated measure one-way ANOVA with Sidak correction (parametric) or repeated measure two-way ANOVA with Sidak’s test (non-parametric) (GraphPad Prism 8.1.0, La Jolla, CA). Data are displayed as mean ±SEM and p values less than 0.05 were considered statistically significant.

## Acknowledgements

This research was supported by NIH NINDS RO1NS076312 and R21NS11945 to LLM and JCC, 1F31NS095568 to LTS, and an Alabama State funded Graduate Research Scholars Program Fellowship (GRSP) to KA. We would like to thank former and current members of the McMahon lab for valuable discussions during the development of this manuscript. This manuscript is hosted as a preprint on biorxiv at https://doi.org/10.1101/672055.

